# Spatial and temporal occurrences of prairie moose across an urban to rural gradient in Saskatoon, Canada

**DOI:** 10.1101/2024.10.19.619215

**Authors:** Kaitlyn E. Harris, Ryan K. Brook

## Abstract

The geographic range of North American moose (*Alces alces*) has expanded over recent decades as the population has recolonized historical habitats and dispersed into new areas, including a notable increase in moose sightings near developed, urban areas. The City of Saskatoon, Saskatchewan is situated on what would traditionally be considered atypical moose habitat, being in the semi-arid, open prairies and surrounded by high-intensity agriculture. However, this area has seen a persistent increase in the frequency of moose occurrences at the urban-rural interface over the last 30 years. We characterized spatial and temporal patterns in moose occurrences over the course of three years (2020-2023) using 29 trail cameras distributed along an urban to rural gradient within the city boundary of Saskatoon. We employed a generalized linear modelling method to assess the potential significance behind where and when moose were occurring along the gradient. Moose occurrence was negatively associated with urban sites containing higher proportions of development (>50% impervious surface cover). While we expected high occurrences during the rut, we found that moose occurrence was low through the fall months. This may be due to the high levels of human disturbance characteristic of the urban-rural interface acting as a deterrent for moose during the breeding season. Future research is warranted to better understand the underlying cause for this result. Moose also occurred most at night, coinciding with the period of lowest visibility and raising concerns for human safety. We provide suggestions and recommendations for future urban moose research and management.

## INTRODUCTION

Moose have long been established primarily as a cold-adapted forest specialist with a broad circumpolar geographic range that incorporates much of northern Canada (Teitelbaum et al. 2021), including the Taiga Plains, Hudson Plains, Boreal Plains, Boreal Shield, Boreal Cordillera, Taiga Shield, and Taiga Cordillera Ecozones (Pastor et al. 1988, Wilken et al. 1996). However, in recent years many moose populations have begun to undergo shifting population trends and changing range dynamics. Timmermann and Rodgers (2017) found that of the 12 Canadian jurisdictions with moose populations, three remained stable (Yukon, British Columbia, Nova Scotia), two were increasing (Quebec, New Brunswick), five were decreasing (Alberta, Saskatchewan, Manitoba, Ontario, Newfoundland), and two had no available estimates (Northwest Territories, Nunavut). Saskatchewan moose populations are among those reported to be declining, and less than half of the number of animals per square kilometer were observed in the 2018 survey of the southern Saskatchewan commercial forest wildlife management zone (WMZ 67) than in the 2004 survey (Government of Saskatchewan 2021).

Despite forage availability being one of the primary limitations to moose populations at the northern extend of their range, warming climates have led to longer growing seasons and increased primary productivity that have contributed to a northern range expansion (Tape et al. 2016, Zhou et al. 2022). At the southern extent, climate change was thought to be a limitation due to temperature-induced heat stress (Wattles et al. 2018a). Thermal heat stress is triggered in moose at temperatures above -5°C in the winter and 14°C in the summer (McCann et al. 2013, Bao et al. 2022). However, while some southern populations have been declining, many others are actively expanding into areas that experience daily ambient temperatures far above the reported tolerance range (Wattles et al. 2018a). This is thought to be explained by behavioural adaptations for reducing thermal heat stress, such as shifting away from the preferential selection of regenerating forest habitats towards forested wetlands and coniferous forest (Wattles et al. 2018a).

Anthropogenic disturbance is also considered as a key driver for ongoing changes to moose distribution (Khan et al. 2022). Certain types of habitat disturbance, such as forestry and logging activities, may positively influence populations through the development of early successional habitats that are favorable for moose (Anderson et al. 2017, Ordiz et al. 2021). Other disturbance features, such as transportation infrastructure, severely limit the distribution of moose and their use of available habitat patches as they have a strong negative response to both high-density road infrastructure and high road use intensity (Wattles et al. 2018b). The risk of moose-vehicle collisions also increases with increasing road density and human activity, and this risk is highest during crepuscular and night times as well as during the seasons associated with increased moose movement (e.g., spring dispersal, fall rut) (Laliberté and St-Laurent 2020). Non-lethal human disturbance, such as off-trail hiking or snowmobile use, will prompt avoidance behaviour in moose although only for a short duration when at a moderate frequency (Neumann et al. 2011). However, disturbance induced by hunting caused no real impact in the activity levels of the general population during the fall moose hunting season, although this effect did vary between individual animals (Neumann et al. 2008).

The potential establishment of prairie moose populations has several important socio-economic implications. Moose feeding activities and trampling have the potential to cause detrimental ecosystem impacts, which can lead to altered vegetation communities (Persson et al. 2000). Furthermore, the substantial overlap in habitat with both wild and domestic ungulates in the prairies has led to an increased risk of disease transmission (Weiskopf et al. 2019). The prevalence of agriculture in this region also means that moose behaviours can be harmful to human livelihood, as they can cause considerable damage to agricultural crops, fence lines, and machinery (Laforge et al. 2017b, Tranulis and Tryland 2023). This may necessitate increased spending on compensation measures for affected landowners (Hanbury-Brown et al. 2021).

Having a contemporary understanding of moose activity on a novel urban prairie landscape is essential for the proper management of this species and for mitigating negative human-moose interactions. We investigated moose occurrences using a network of wildlife monitoring trail cameras distributed within the city boundary of Saskatoon, Saskatchewan, Canada. Our objectives were to (1) investigate the spatial patterns of moose occurrence relative to the proportion of impervious surface cover (e.g., roads, buildings) along an urban to rural gradient, and (2) identify any temporal trends in occurrence by analyzing the baseline annual, monthly, and daily moose occurrences throughout the study period. We further aim to use these data to assess the potential for a resident prairie population of moose in Saskatchewan. We expected moose occurrences to be associated primarily with areas of low development density (i.e., the rural end of the gradient) (Tinoco Torres et al. 2011, Laforge et al. 2016). We predicted moose occurrence in Saskatoon would be low through the winter and during the calving season (Eriksen et al. 2011), and that moose would occur most mid-summer (following patterns in plant phenology) (Van Ballenberghe and Miquelle 1990) and during the fall rutting period (Philips et al. 1973). We further hypothesized that moose occurrence would be highest at night, as many species near urban boundaries respond to anthropogenic disturbances by shifting to primarily nocturnal activity patterns (Gaynor et al. 2018, Procko et al. 2023).

## STUDY AREA

This study took place within the city boundary of Saskatoon, Saskatchewan, Canada (Fig. 1), which has an urban population of 266,000 residents and a land area of 227 km^2^ (Statistics Canada 2021). Moose sightings were rare in the semi-arid Prairie Ecozone of Western Canada until approximately three decades ago, when their range appeared to expand southward from the northern boreal forest and southern boreal-deciduous fringe to now broadly include the Canadian prairies (Laforge et al. 2017b). Saskatchewan, the central prairie province, is one of the largest worldwide exporters of agricultural crops, containing approximately 24 million hectares of total farm area, 67% of which is designated cropland (St. Pierre and Mhlanga 2022). This agriculture-dominated landscape is included in the Prairie Pothole Region (PPR), and contains many thousands of wetland potholes embedded in a complex agro ecosystem dominated by annual and perennial crop cover (Pennock et al. 2010).

**Fig. 1.**
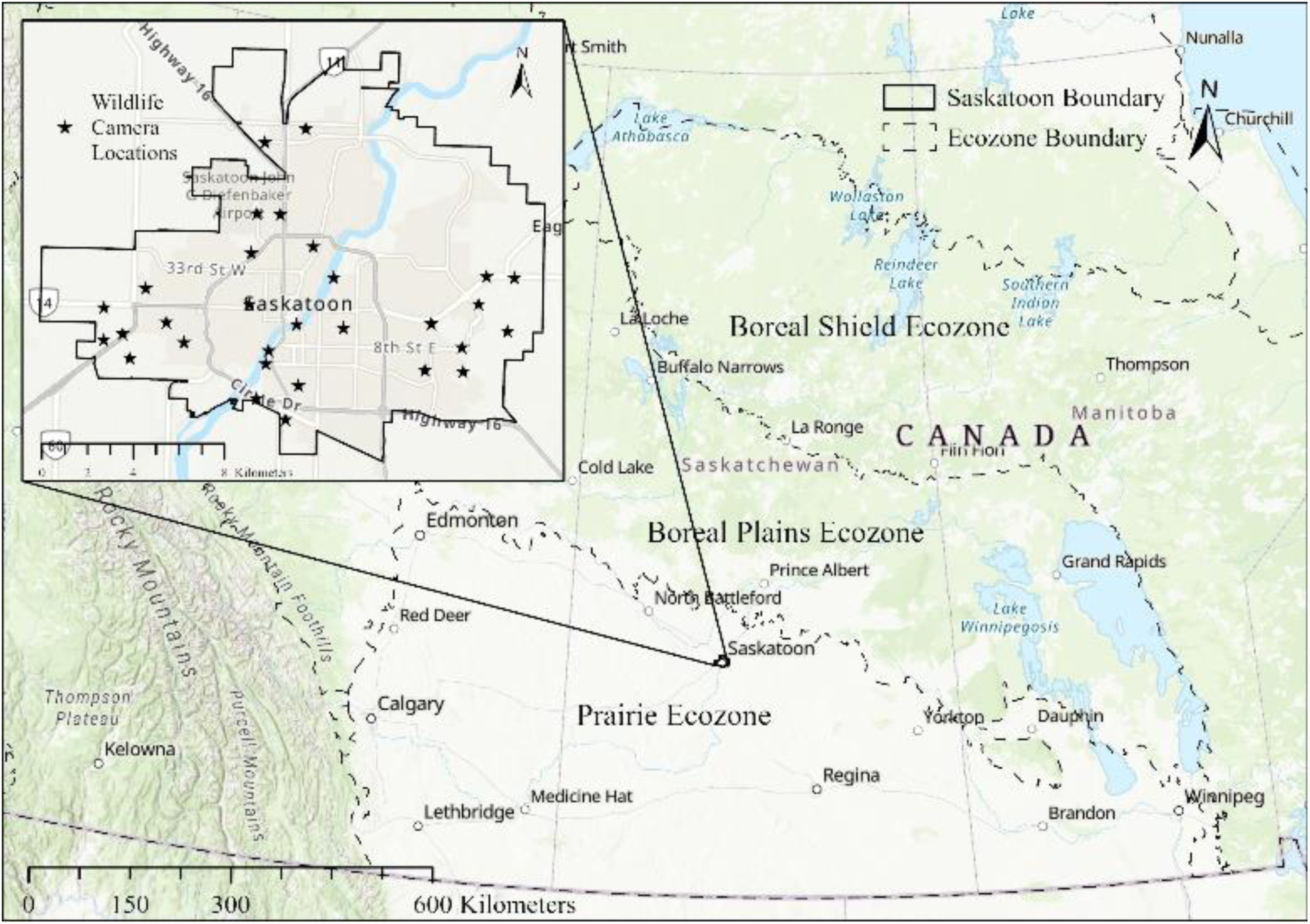
The location of the study area relative to the Boreal Shield, Boreal Plains, and Prairie ecozones, and the distribution of all wildlife monitoring trail cameras (n=30) deployed in a stratified random distribution along an urban to rural gradient within the city boundary of Saskatoon, Saskatchewan, Canada.

## METHODS

The provincial research permit for this study was issued by the Government of Saskatchewan’s Ministry of Environment (Academic Research Permit Number: 20AR010w) and was maintained through quarterly dataset submissions. The camera trap locations were pre-determined using ArcGIS software (v. 10.7.1, ESRI Inc. 2019) in accordance with protocols provided to member cities by the Urban Wildlife Information Network (UWIN) (Magle et al. 2019). The City of Saskatoon was digitally stratified into six primary land cover classifications (agricultural, aquatic, built (i.e., impervious surface cover), forested and shrubland, grassland, and greenspace) using the Meewasin Valley Authority Natural Areas Inventory land cover map (Hooey 2021).

Two perpendicular transects were created based on the four cardinal directions (North-South and East-West) and layered over the Saskatoon land cover map, originating from a central downtown location (52°7’24” N 106°39’50” W) and extending outwards to the city boundary. The North-South transect measured 16 km in length and the East-West transect measured 20 km in length. Stratified random sampling (n=30 samples) randomly placed five points within each of the six land cover classifications using spacing parameters of a 1 km minimum distance between points, and points fixed within 1 km of either side of the transect. This design helped to ensure the independence of data points as well as promoted representative sampling of potential urban habitats.

Once the points were distributed, a 500 m radius buffer was run around each point and used to quantify the proportions of each identified land cover within that zone. Any areas with incomplete coverage were assigned an initial value of “null” and visually quantified using City of Saskatoon orthoimagery. The landcover classification analysis was used to define the urban to rural gradient by classifying each buffer zone in ArcGIS based on the “built” landcover class, which identified the proportions of impervious surface area within each zone. Impervious surface area referred to the proportion of developed surface cover in an area (e.g., roads, buildings, parking lots). Rural zones contained <25% impervious surface area, peri-urban zones contained 25-50% impervious surface area, and urban zones contained >50% impervious surface area. The final gradient consisted of nine cameras (30%) within rural areas, nine cameras (30%) within peri-urban areas, and 12 cameras (40%) within urban areas. Four potential camera trap locations in each zone were selected based on accessibility and suitability of the location for year-round monitoring, and the final camera positions were determined following ground truthing and landowner agreements.

### Camera Traps

The trail cameras (n=30) in this study were Browning Dark Ops HD Pro X motion-activated infrared trail cameras. Each camera trap consisted of a trail camera (containing six lithium AA batteries and one 128GB SD card) padlocked inside a Browning security box and secured to a tree using a tree strap, a Python adjustable cable lock, and a length of 5/16” transport chain. The cameras were left on default settings (trail mode with high detectability) and set to take a burst of three images at 1 second intervals with each activation and a 30 second delay between activations. To reduce the potential effects of autocorrelation in trail camera data, individual wildlife occurrences were comprised of a photo event consisting of 1+ images of an animal taken at a single location in time and space (i.e., a single camera trap). Therefore, a single wildlife occurrence in this study was defined as any photo event of the same animal captured by the same camera trap within a 60 min timeframe (Leblond et al. 2007, Laforge et al. 2017a).

### Data Collection

Trail camera deployment occurred on 8 September 2020 and this first phase of the project ended 9 September 2023. Each camera trap was positioned 1.5-2 m above the ground and, where possible, faced a natural corridor or trail with minimal vegetation in front to minimize accidental triggering by moving vegetation. The camera traps ran continuously, year-round and were checked at regular intervals by research staff. Camera cards were pulled and replaced at every visit and camera servicing, maintenance, and replacements were performed as necessary. All wildlife photos were uploaded to an online database provided to UWIN member cities (urbanwildlifenetwork.org) and all human data was discarded prior to image uploading, as stipulated in the landowner agreements. Each image was individually examined and tagged in the database. Greatly distorted or unclear images were tagged as “unknown.” Each photo tag was automatically assigned the associated metadata (i.e., image time and data, camera number, site identifier, camera GPS location coordinates), and was accessible for download directly from the database. At each site we also estimated the green canopy cover (%) above each camera using the Canopeo application (Patrignani and Ochsner 2015), and the average height of vegetation, which was calculated based on field measurements of the highest point of herbaceous vegetation below each camera and 10 m in front of each camera. One camera trap (camera #19) was removed from the final data analysis due to chronic vandalism and theft that created major gaps in the dataset.

Moose occurrences were summarized and sorted by year, month, time-of-day, and dominant landcover type. The time-of-day classification was standardized based on hours of daylight to account for the annual fluctuations in Saskatoon daylength. Hours of daylight were calculated as the total difference in hours between the true sunrise and sunset times (obtained from: timeanddate.com) for each unique occurrence date, and each moose occurrence was categorized based on when the image was captured relative to the calculated daylight hours. Any occurrence within the first 25% of daylight hours was considered morning, within the following 50% of daylight hours were considered afternoon, and within the final 25% of daylight hours were considered evening. Any occurrence after sunset or before sunrise was considered night. The dominant landcover type was based on the ArcGIS landcover classification analysis, whereby the landcover class found to be in the highest proportion within each buffer zone was designated as the dominant type for that site. Where possible, occurrences were further classified by the age and sex of the animal by visually examining each photo and using key identifier characteristics. Moose were considered adults at one year of age. Young calves were identified by smaller body size, lighter or reddish colored fur, small nose and short ears, and very small to no bells. Adult females were identified based on a more consistent and lighter colored face, small bells, and a white vulva patch beneath the tail. Bull moose were identified based on a much darker nose, large bell, no vulva patch, and presence of antlers.

### Statistical Analysis

We employed a generalized linear modelling (GLM) approach to assess the potential significance of when and where moose were occurring along the urban to rural gradient. Our initial model was fit with a poisson distribution and contained temporal predictor variables (month and time-of-day) as well as habitat data (green canopy cover, average vegetation height, dominant landcover class, and the gradient class of each site). An interaction between month and time-of-day was also included to assess their combined potential influence on moose occurrence. We adjusted our model to a quasi-likelihood approach after calculating the dispersion parameter of the poisson model and finding slight over-dispersion in the model residuals (dispersion > 1) (Ver Hoef and Boveng 2007). Because Akaike’s Information Criterion (AIC) is undefined for quasi-poisson models, we employed a stepwise selection process to identify the best fit model through model comparison and testing the removal of non-significant predictors one at a time using a likelihood ratio (Chi-square) test. The best fit model was selected once we reached a base model where the removal of any predictor variable showed a significant loss of explanatory power. We also performed a Kruskal-Wallis Test to assess whether moose occurrence differed significantly between the three study years. All statistical analyses were performed using R statistical software (v. 4.2.1, R Core Team 2022), and an alpha level of *p* ≤ 0.05 was used to identify statistical significance.

It must be noted that one study site accounted for over half of the observed moose occurrences. To explore the potential influence of this site on our overall results, we conducted a supplementary analysis by re-running the GLM model selection process with this site excluded from the dataset. This approach allowed us to evaluate whether the observed effects were relatively consistent across the broader study area or being driven primarily by this site.

## RESULTS

There were 60 moose occurrences at 12 of 29 sites over 30,230 camera trap days (Fig. 2). While our results indicated no statistical difference between study years, the total moose occurrence did increase each year over three years (Sept 2020 – Sept 2021: n=8, Sept 2021 – Sept 2022: n=23, Sept 2022 – Sept 2023: n= 29). Adult cows occurred in the highest overall frequency during the study period (n=18), followed by adult bulls (n=14), and cow-calf pairs (n=5 pairs). Cow-calf pairs occurred yearly between March – July (2021: n=1 pair, 2022: n=2 pairs, 2023: n=2 pairs). There was also n=4 occurrences that were classified as “calves of the year (COY).” The COY group included occurrences of young calves detected between July – October. This group was excluded from the cow-calf pair grouping as no adult was present in the images, although it is assumed that the mother simply remained out of the camera line of sight. There was n=14 moose occurrences that we were unable to classify by age or sex due to poor image quality.

**Fig. 2.**
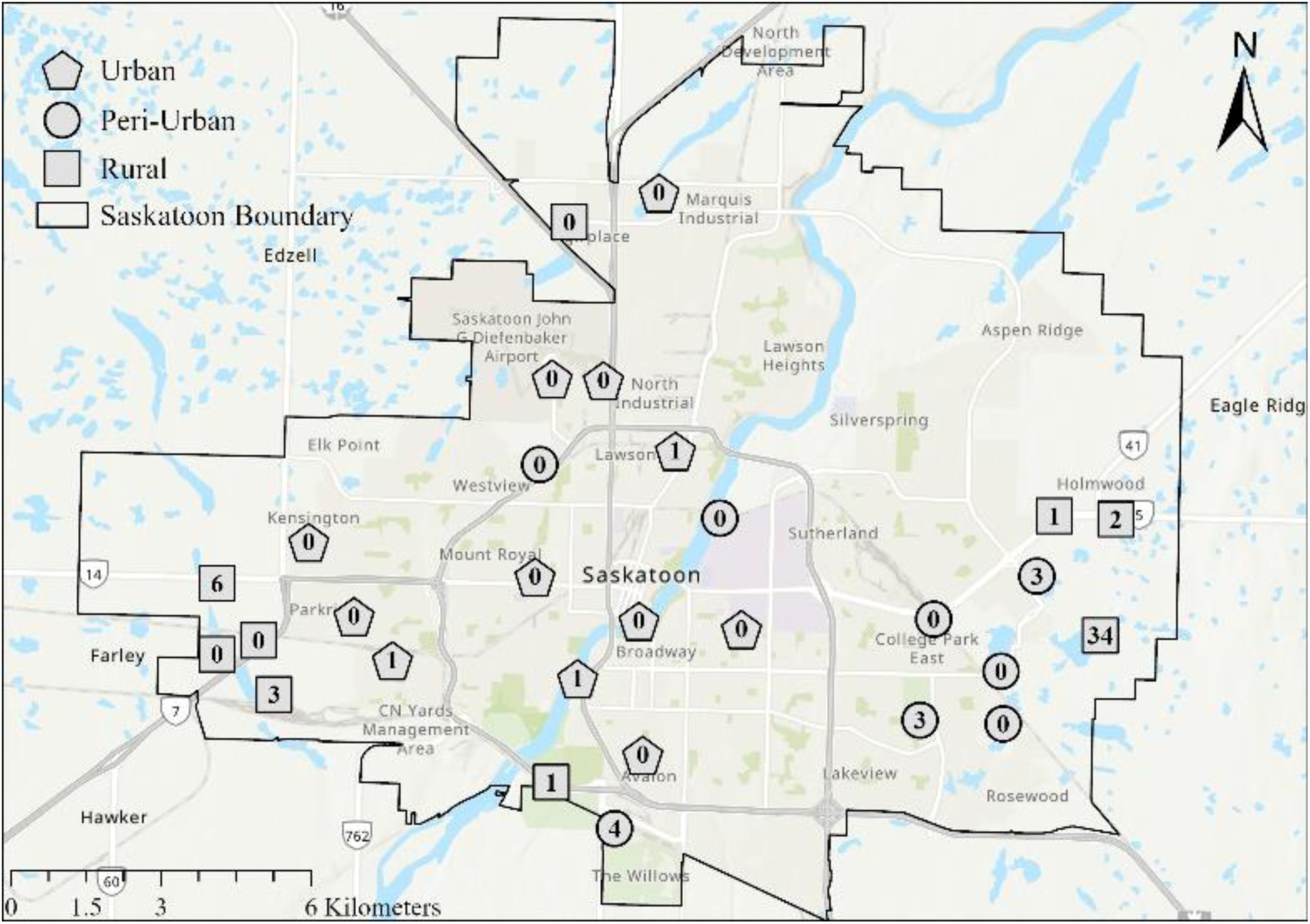
Total (n=60) moose occurrences per camera trap obtained from trail cameras (n=29) deployed in a stratified random distribution along an urban to rural gradient within the city boundary of Saskatoon, Saskatchewan, Canada, 2020-2023.

### Moose Occurrence Model

The model selection process did not support the inclusion of a temporal interaction between month and time-of-day in the base model (*p*=0.41). We also investigated the addition of vegetation height and green canopy cover as potential predictors for moose occurrence, however the model comparison results indicated no significant loss of explanatory power with their exclusion from the base model (*p*>0.05, Table 1). The best fit moose occurrence model contained the variables for month, time-of-day, gradient class, and dominant landcover class (*p*<0.05, Table 2).

**Table 1.** GLM model comparison results for selecting the best fit model of moose occurrence in Saskatoon, Saskatchewan, Canada, 2020-2023. Number of model variables (K) and Chi-square model comparison score (Chi). A significant Chi-square comparison score indicated a significant loss of explanatory power after removal of a model variable.

| Model | K | Chi |
| --- | --- | --- |
| Month * Time_of_Day + Gradient + Avg_Veg_Height +<br>Canopy_Cover + Dom_Landcover | 6 | NA |
| Month + Time_of_Day + Gradient + Avg_Veg_Height +<br>Canopy_Cover + Dom_Landcover | 6 | 0.41 |
| Month + Time_of_Day + Gradient + Canopy_Cover + Dom_Landcover | 5 | 0.67 |
| **Month + Time_of_Day + Gradient + Dom_Landcover** | 4 | 0.34 |
| Month + Time_of_Day + Gradient | 3 | 0.011* |
| Time_of_Day + Gradient + Dom_Landcover | 3 | 0.017* |
| Month + Time_of_Day + Dom_Landcover | 3 | 0.0086** |
| Month + Gradient + Dom_Landcover | 3 | <0.001*** |

**Note:** \* $p < 0.05$ , \*\* $p < 0.01$ , \*\*\* $p < 0.001$

**Table 2.** Chi-square analysis of the best moose occurrence model based on temporal and habitat data along an urban to rural gradient in Saskatoon, Saskatchewan, Canada, 2020-2023.

| **Month + Time_of_Day + Gradient + Dom_Landcover** |  |  |  |
| --- | --- | --- | --- |
| Model variable | Df | Deviance resid. | Pr(>Chi) |
| Month | 11 | 37.99 | 0.0011** |
| Time-of-day | 3 | 48.48 | <0.001*** |
| Gradient class | 2 | 64.27 | <0.001*** |
| Dominant landcover class | 4 | 16.01 | 0.011* |

**Note:** \* $p < 0.05$ , \*\* $p < 0.01$ , \*\*\* $p < 0.001$

One rural site located at the eastern edge of the study area observed 57% of the total moose occurrences, and the exclusion of this site did have some influence on our model selection results. The month and dominant landcover class variables lost significance in the base model, while the time-of-day and gradient class remained as the primary predictor variables for the best moose occurrence model.

### Spatial Moose Occurrence

Moose occurred predominantly along the eastern edge of Saskatoon, although they also occurred along the western and southern edges of the city, all of which are zoned primarily as residential, and at several interior sites adjacent to the river corridor (Fig. 2). No moose were captured by camera traps in the northern, industrial zoned section of Saskatoon. While moose did occur in all categories of the urban to rural gradient, they primarily selected against urban sites containing greater than 50% impervious surface cover (*p*<0.05, Fig. 3).

**Fig. 3.**
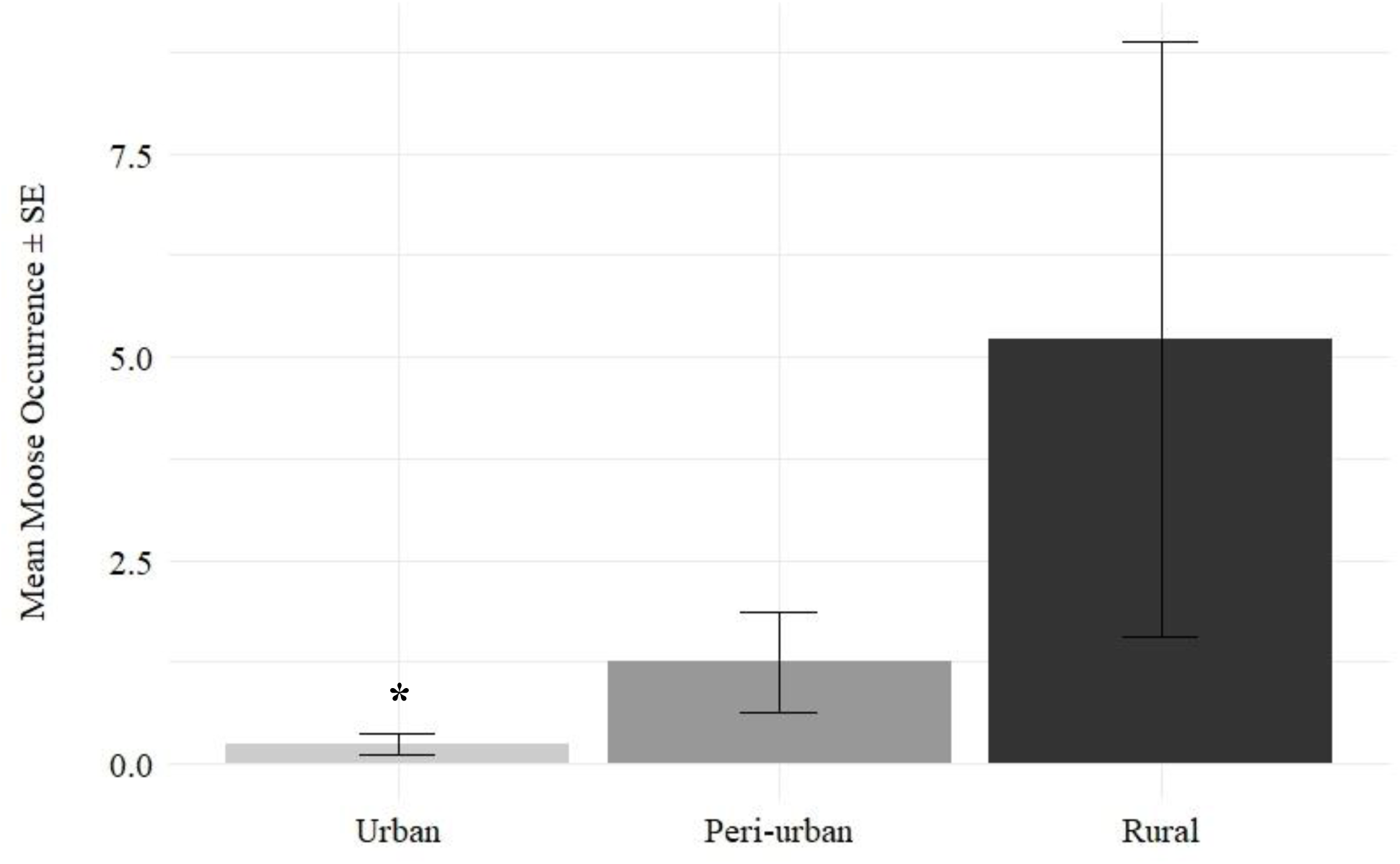
Mean and standard error (SE) of moose occurrences within urban, peri-urban, and rural sites in Saskatoon, Saskatchewan, Canada, 2020-2023. Urban sites contained <25% impervious surface area, peri-urban sites contained 25-50%, and rural sites contained <25%. Asterix (*) indicates a significant statistical difference between variable levels based on best fit model results (*p*<0.05).

While the dominant landcover class was found to be a significant predictor in the best fit model, there were no statistical differences found between any pairs of landcover classes. This is likely due to the limited variation within our study area, as most of the 29 camera sites were dominated by the “built” landcover class (n=17). This was followed by “agriculture” (n=8), “greenspace” (n=2), and then “aquatic” and “grassland” (n=1, respectively). No sites had “forest and shrub” as its dominant landcover class. While trees and/or shrubs were present at all sites, only four sites contained >7% total forest and shrub cover, and most sites contained <2%.

### Temporal Moose Occurrence

We observed some variation in seasonal occurrence patterns. Contrary to general knowledge, moose occurrence in our study area was low throughout the fall months (September – November), and had a significant negative association with the month of October (*p*<0.05, Fig. 4). These months are associated with the rutting season and, typically, high moose activity. Mean monthly occurrences remained mostly low through the winter months before increasing in April, dropping in May (the calving season), and then peaking in July. February was the only month without moose occurrences across all years. The time-of-day also had a significant influence on moose occurrence. Moose occurred most frequently at night (*p*<0.05, Fig. 5), indicating an aversion to human activity.

**Fig. 4.**
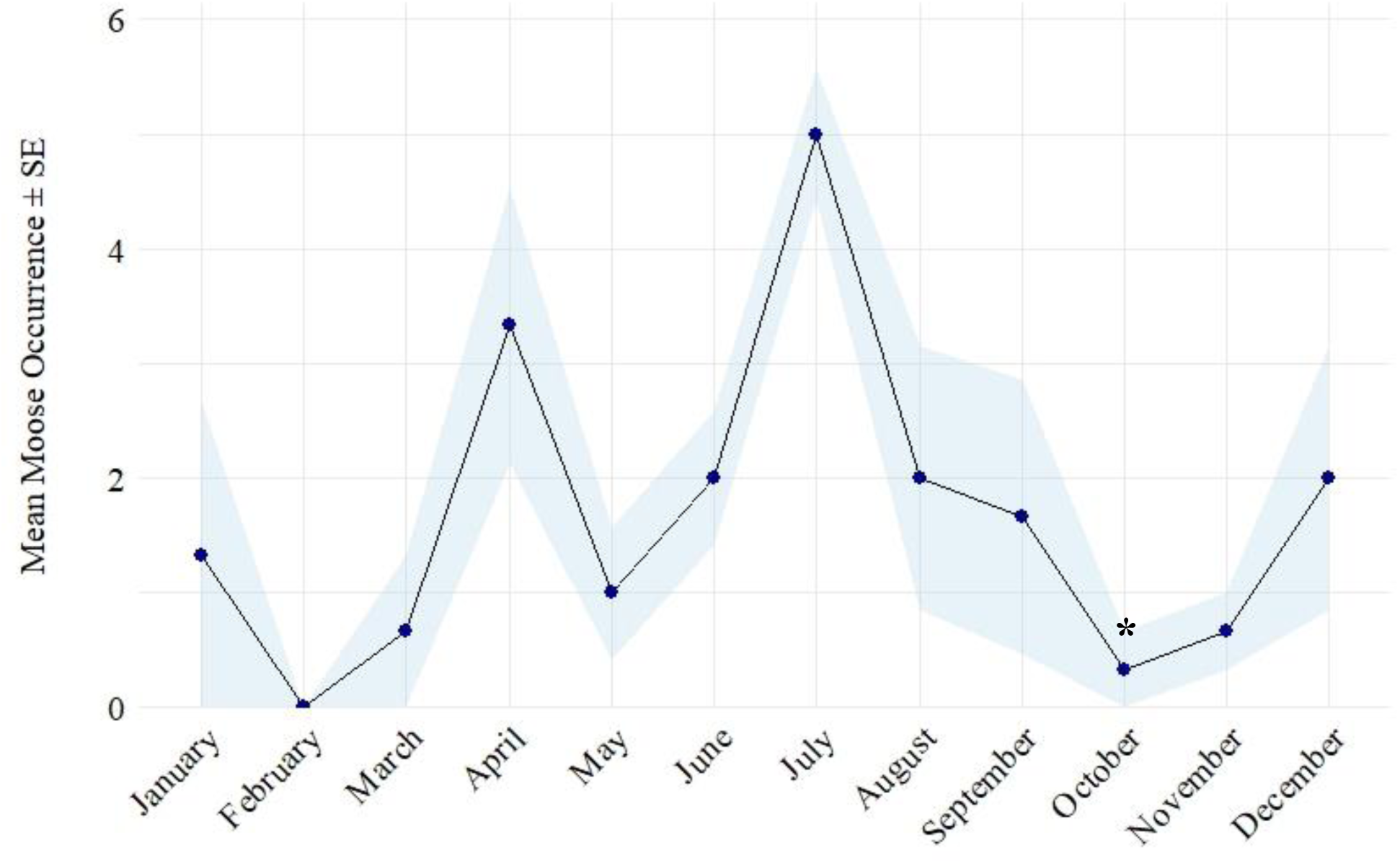
Mean and standard error (SE) of moose occurrences per month in Saskatoon, Saskatchewan, Canada, 2020-2023. Asterix (*) indicates a significant statistical difference between variable levels based on best fit model results (*p*<0.05).

**Fig. 5.**
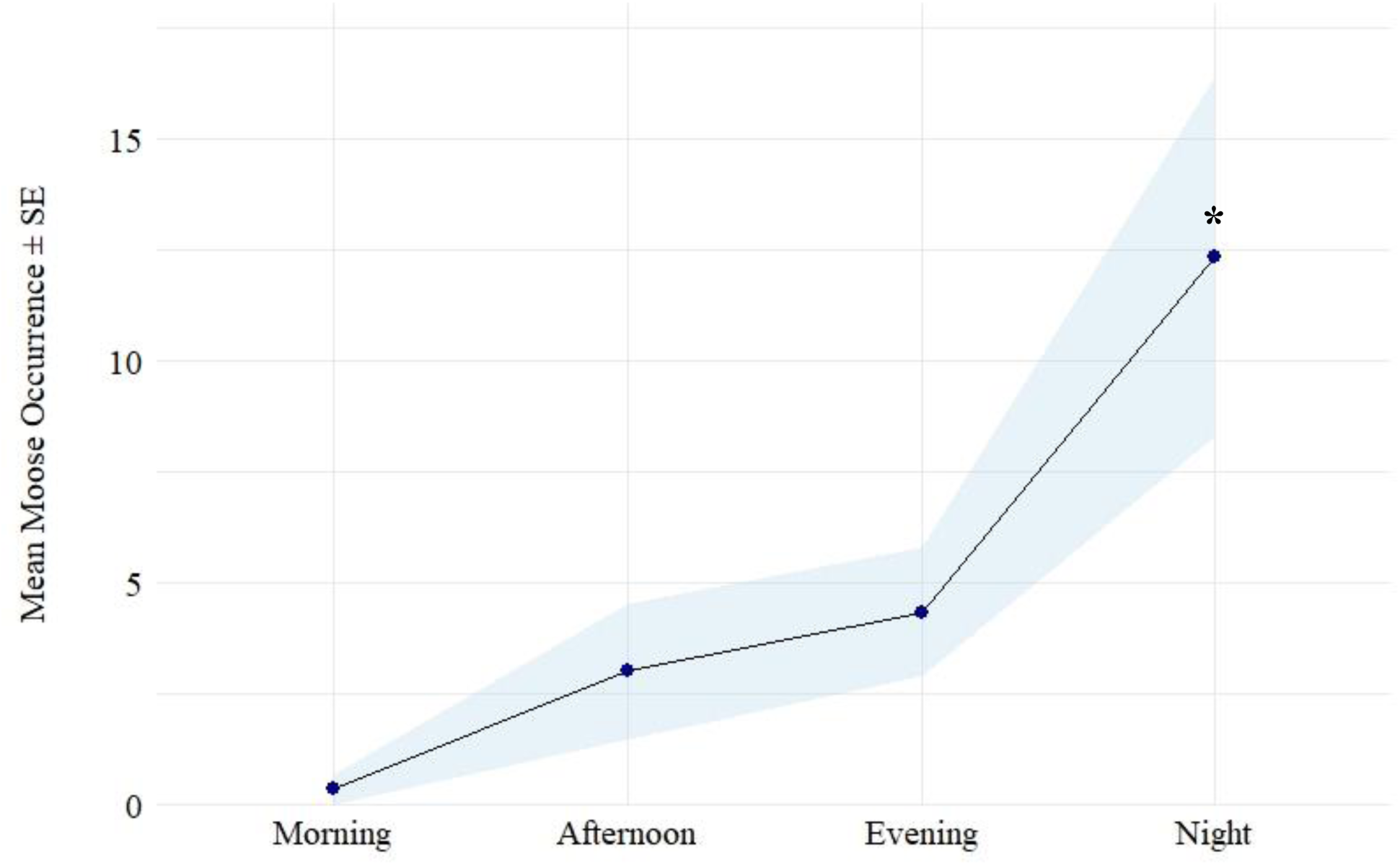
Mean and standard error (SE) of moose occurrences per time-of-day in Saskatoon, Saskatchewan, Canada, 2020-2023. Asterix (*) indicates a significant statistical difference between variable levels based on best fit model results (*p*<0.05).

## DISCUSSION

Moose have long been considered as a forest specialist species, thought unable to thrive in the Canadian prairies due to the overly warm, semi-arid climate, lack of forest cover, and landscape composition (Laforge et al. 2016). In particular, the distinct lack of vertical hiding and thermal cover on the prairies, where summer temperatures can reach extremes of >35°C, creates a significant risk for thermal heat stress in moose. However, the widespread distribution of small, tree-ringed wetlands found throughout the PPR in tandem with the abundant croplands of the prairies appears to have created a novel environment for moose to exploit. These wetland sites allow moose to thermoregulate through submersion while having access to the surrounding high-quality farm crops as a supplementary food source (Laforge et al. 2016). We believe these landscape factors are a key component to the ongoing moose population expansion in the prairies.

Alongside the documented range expansion is an increase in moose occurrences near sub-optimal urban landscapes. These include, aside from Saskatchewan, rising populations in proximity to large urban settlements in regions such as Alaska (Welch et al. 2015), Utah (Wolfe et al. 2010), Massachusetts, New York, and Connecticut among others (Wattles and DeStefano 2011). While it is less common for moose to enter the developed city proper, they are demonstrating adaptability to near-urban habitats. This may be a result of the higher levels of primary productivity from anthropogenic inputs relative to the surrounding area acting as an attractant (Welch et al. 2015). The common use of road salt to de-ice winter roads, leading to the formation of roadside mineral sources, may also be a contributing factor (Leblond et al. 2007, Rea et al. 2021). Regardless, the dense human populations, dense road infrastructure, and high road use intensity in urbanized habitats makes the establishment of moose populations in these areas hazardous to human safety.

Our results indicate that moose occurrence, while increasing annually, did vary throughout the year. Occurrences remained low through the winter months before increasing in April. This is likely due to post-winter dispersal associated with warming spring temperatures, melting snow, and increased forage distribution and quality (Risenhoover 1986, Laliberté and St-Laurent 2020). The displacement of yearling calves prior to the new calving season may further be contributing to higher springtime occurrences (Testa 2004). Moose occurrences dropped low during the calving season (May), before peaking in July. This is consistent with other findings detailing the reduction in moose activity while calving (Eriksen et al. 2011), and the subsequent increase in activity post-calving and through the summer months that follows patterns in plant phenology (Van Ballenberghe and Miquelle 1990). However, contrary to both general knowledge and our predictions, we found low fall occurrences in our study area and a significant negative association with the month of October. Fall activity is influenced by the rut, with high daytime activity and the aggregation of social groups that are not seen at other times of the year (Best et al. 1978, Luymes et al. 2024). The significant lack of moose activity that we observed during this period is notable, as we also found evidence of a resident breeding population. This was based on the yearly presence of young calves and cow-calf pairs detected during and following the calving season, as well as the year-round occurrences of adults of both sexes.

Vocalizations play a key role in moose mating behaviour, serving as a means for finding and attracting potential mates (Bowyer et al. 2020). While a minor to moderate amount of human disturbance may not significantly influence fall moose activity (Neumann et al. 2008), the high levels of human activity and disturbance characteristic of the urban-rural interface may be acting as a deterrent for moose during the breeding season. Further research on seasonal moose activity in urbanized areas is warranted to better understand the mechanism underlying this finding.

Daily occurrence patterns were as expected, with occurrences lowest during the daylight hours and increasing in the evening before being significantly highest at night. This is mostly consistent with the known activity patterns of moose being most active at crepuscular and night times (Risenhoover 1986, Dussault et al. 2006). The strong nocturnal preference and avoidance of highly developed sites in our study area suggest that moose may be demonstrating an anthropogenic avoidance behaviour through a temporal adaptation, which is common in many wildlife species inhabiting areas with high levels of human activity and disturbance (Procko et al. 2023). Since moose-vehicle collisions tend to occur in the highest frequencies at dawn, dusk, and night (Dussault et al. 2006, Klassen and Rea 2008), characterizing the temporal activity patterns of moose inhabiting urban-rural transition habitats is essential to minimize the risk potential, especially during the periods of low visibility when moose are most active.

### Management Implications

Moose are not compatible with urban environments due to their large size, potential for aggression, and the significant safety risks associated with negative human-moose interactions. Key risks that require further consideration include the potential for property damage (e.g., gardens, fences, trees, shrubs, etc.), safety issues related to moose injuring humans due to aggressive behaviour, moose-vehicle collisions, and moose-train collisions. Management preparation and response considerations should include increasing public awareness and education, installing fencing to keep moose out of areas that are of highest concern as well as one-way fencing that will allow moose to escape while preventing them from coming back in, improved signage that is larger and lighted to better warn drivers in high-risk areas, and continued support of wildlife immobilization and translocation programs. Understanding moose activity and occurrence trends in a novel environment requires consistent and long-term monitoring. Future research should, if possible, incorporate a greater number of wildlife monitoring camera traps, investigate the seasonal and temporal patterns identified in this study with respect to annual variations in human activity and environmental conditions, further examine population dynamics, and analyze and evaluate entry points and hotspots of greatest risk.

## ACKNOWLEDGEMENTS

Our study was supported by the Natural Sciences and Engineering Research Council of Canada (NSERC), the Urban Wildlife Information Network (UWIN), the University of Saskatchewan, the College of Agriculture and Bioresources, and the Department of Animal and Poultry Science. We thank our partners at Meewasin Valley Authority, Wild About Saskatoon, the Saskatoon Forestry Farm Park and Zoo, the City of Saskatoon, and the Saskatoon Parks Department. We also thank Giselle Hooey, Candace Savage, Renny Grilz, and Douglas Clark for their individual contributions.

## Notes

### Competing Interest Statement

The authors have declared no competing interest.

